# Fast and artifact-free excitation multiplexing using synchronized image scanning

**DOI:** 10.1101/2023.02.07.527342

**Authors:** Ezra Bruggeman, Robin Van den Eynde, Baptiste Amouroux, Tom Venneman, Pieter Vanden Berghe, Marcel Müller, Wim Vandenberg, Peter Dedecker

**Affiliations:** Department of Chemistry, KU Leuven, Belgium; Yusuf Hamied Department of Chemistry, University of Cambridge, UK; Department of Chronic Diseases Metabolism and Ageing, KU Leuven, Belgium

## Abstract

We present the Resonator, a simple optical device that provides quasi-simultaneous fluorescence imaging with multiple excitation wavelengths. The device uses a resonant scanning mirror to periodically displace the sample image on a camera sensor at a rate that is much faster than the image acquisition rate. The excitation light is synchronized with the scanner motion to create two laterally shifted copies of the image, each containing the fluorescence excited by a single wavelength. The additional information is then encoded either into the point-spread function of the imaging or as multiple distinct images. Since this multiplexing is performed at very high rates, our design can eliminate or mitigate artifacts caused by temporal aliasing in conventional sequential imaging. We demonstrate the use of our system for the monitoring of fast light-induced dynamics in single quantum dots and for the imaging of Ca^2+^ signalling in hippocampal neurons.

## Introduction

Multicolor fluorescence microscopy is a workhorse technique in the life and materials sciences. To prevent distortions and artifacts, the different color channels must be acquired in a timespan that is short compared to the sample dynamics. In many experiments sequential (‘one by one’) acquisition of the different channels can work well, provided that these dynamics occur on a timescale of tens to hundreds of milliseconds or slower. However, sequential imaging can readily lead to temporal aliasing artifacts when faster dynamics are encountered.

Avoiding such artifacts is best done by acquiring the different color channels simultaneously. This is conceptually easy to do when the channels differ in emission spectrum, since these can be straightforwardly split using dichroic mirrors and optical filters. Readily available solutions include the use of image splitters that focus the different emission bands onto distinct regions of the camera sensor. In sufficiently sparse samples, information on the emission color may also be obtained in other ways, such as the direct analysis of the shape of the point-spread function (PSF) [1], possibly enhanced by PSF engineering [2–5], or the introduction of a dispersive optical element such as a prism [6, 7] or diffraction grating [8–10]. Overall, the simultaneous acquisitions of multiple emission bands usually works well because the full required information (the wavelength of the light) is contained within the emission photons themselves.

These solutions stand in stark contrast to the much more limited possibilities for simultaneous multicolor imaging based solely on the excitation spectra of the fluorophores. This need is encountered in a variety of situations, such as the use of excitation-ratiometric biosensors [11, 12], (single-molecule) FRET [13, 14], single-molecule localization microscopy [15], or when studying the spectroscopy of complex photochemical systems such as quantum dots [16]. However, (near-) simultaneous multiplexing based on excitation wavelength is a much more difficult challenge because an emission photon does not ‘know’ the excitation wavelength that caused it to be emitted. As a result, sequential acquisitions remain the gold standard and temporal aliasing issues can readily arise except in select cases where the information content is limited. A number of alternative strategies have been explored, such as the encoding the excitation wavelength in the temporal modulation of the light sources [15, 17, 18] or stroboscopic illumination [19]. However, these methods likewise induce a temporal delay in the imaging.

In this work, we develop near-simultaneous excitation multiplexing by synchronizing the illumination with rapid displacements of the sample image on a camera sensor. This strategy can take advantage of the availability of fast and inexpensive displacing elements such as resonant scanners or micro-mirror devices that easily provide scanning frequenties of tens of kHz, as well as the microsecond or faster modulation allowed by even inexpensive light sources. We realize this idea via a proof-of-concept implementation called the ‘Resonator’, an optical module that can be straightforwardly added onto existing widefield fluorescence microscopes, and show that it can deliver near-simultaneous multiplexing of two different excitation wavelengths. Depending on the optical configuration and scan settings, this device can be used to encode the spectral information into the point-spread function (PSF) of the imaging, or in a more classical arrangement where the different excitation wavelengths are imaged onto distinct regions of a camera. We explore some of the possibilities enabled by this device by imaging fast dynamic processes occurring in single quantum dots as well as a biosensor reporting calcium dynamics in hippocampal neurons.

## Results and discussion

### Optical design

The overall design of our system is shown in figure 1a. The key feature is the introduction of a resonant scanner into the emission beampath, which induces fast and periodic image displacements on the camera sensor. Such scanners are broadband mirrors that continuously and reliably oscillate at a fixed frequency up to tens of kHz depending on their construction, and that are furthermore available at modest expense. To complete the optical pipeline, we also introduced two additional lenses: a lens that collimates the light focused by the microscope tube lens, and a second lens that focuses the scanned light onto the camera.

**Figure 1:**
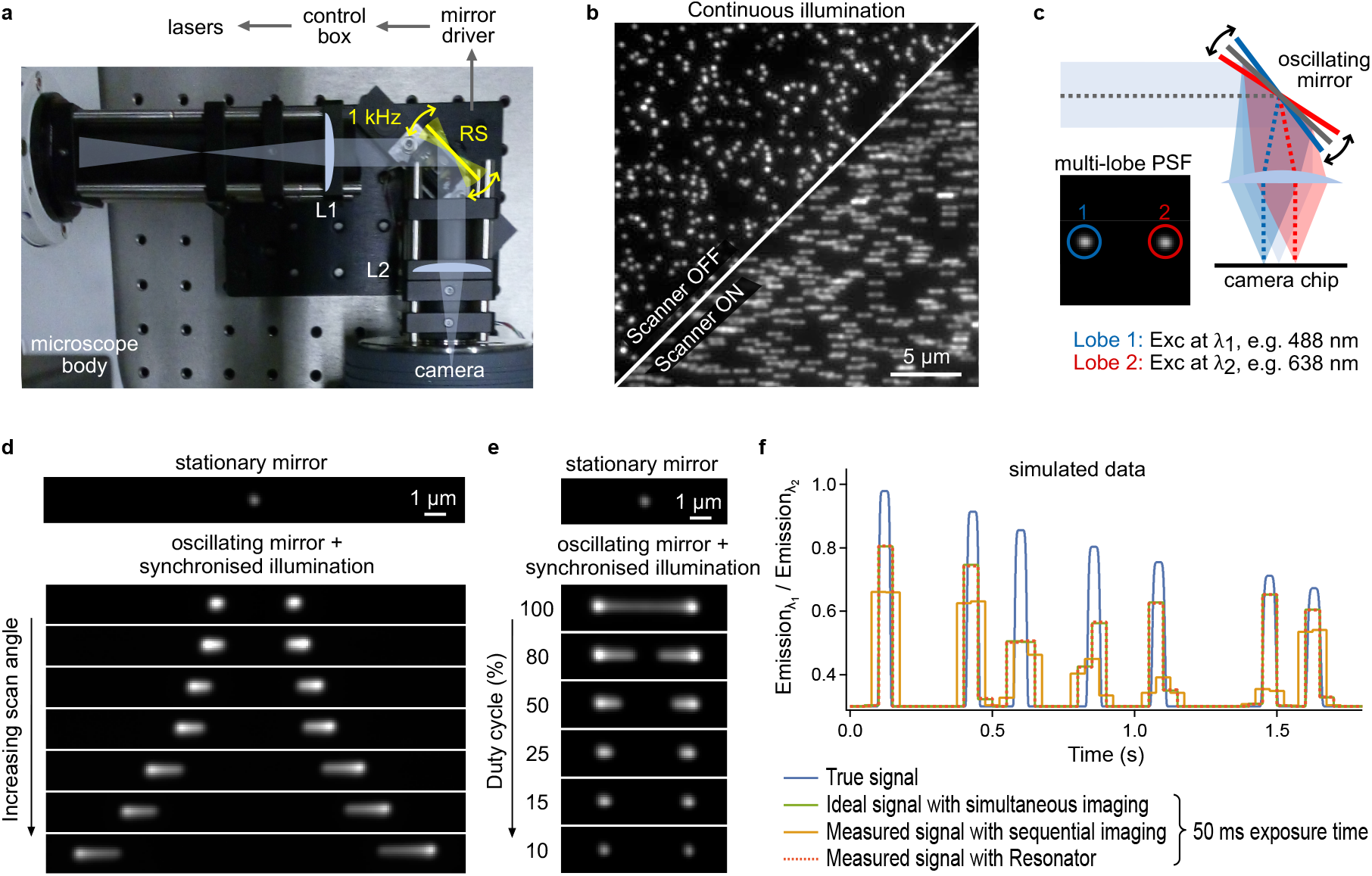
Concept. **(a)** Optical layout of the design including two relay lenses (L1 and L2) and the resonant scanner (RS) that displaces the image on the camera at 1 kHz. **(b)** Image of fluorescent beads (0.1 µm Tetraspeck beads) with and without oscillating motion of the scanner and 100% illumination duty cycle (i.e. laser is on continuously). **(c)** Schematic illustration showing the synchronization of the illumination with the scanner motion. **(d)** Image of a single fluorescent bead showing the response of the system for different scanning amplitudes. **(e)** Image of a single fluorescent bead showing the response of the system for different duty cycles of the illumination. **(f)** Simulated data showing the (near-) elimination of aliasing artifacts using the Resonator. The blue line shows simulated ground-truth data resembling the oscillatory response of an excitation-ratiometric biosensor. The red trace shows the ideal but experimentally infeasible responses for true simultaneous measurements using a finite camera exposure time. The green and orange traces show the expected responses for sequential imaging and the use of our Resonator with a resonant scanner running at 1 kHz. Higher frequencies of the resonant scanning render the signal indistinguishable from the ideal (green) trace.

Activating the resonant scanner results in the formation of a ‘smeared-out’ image under continuous illumination (figure 1b), reflecting the fact that the scanning frequency is much faster than the camera acquisition rate, which is usually in the tens of milliseconds or longer regime. This smearing can be avoided by synchronizing the illumination with the motion of the scanner, applying one illumination wavelength when the mirror is close to one of its turnaround points, and a second illumination wavelength when the mirror is close to the other turnaround point (figure 1c). The net result is that a single emitter appears as two different spots in the image, where the distance between the spots is governed by the scan amplitude (figure 1d) and their shape is determined by the duty cycle (figure 1e), defined here as the percentage of the time that at least one of the light sources is active. If the illumination duration can be made sufficiently short by using high-intensity light sources, more excitation wavelengths can furthermore be introduced at additional points of the scanning cycle. The relative brightness of the spots is determined by the relative light intensities and the efficiency with which the emitter can be excited by the different wavelengths. Our device thus provides an estimate of the excitation efficiencies for every emitter within the sample.

A crucial advantage of the Resonator is the fast operation enabled by the resonant scanner. Using a scanner operating at 16 kHz, for example, the time to switch between excitation wavelengths is only 31 µs, which is orders of magnitude faster than conventional sequential acquisitions, and also independent of the rate at which images are acquired since the signals are simply integrated by the detector. While the device does not increase the overall image acquisition speed, and hence in principle does not allow more frequency content to be acquired, it does strongly mitigate the possibility of aliasing artifacts since both excitation wavelengths are probed very rapidly and over the full duration of the acquisition, as we show using simulated data in figure 1f. In this figure and what follows, we denote the emission excited by irradiating using a wavelength *λ* as ‘emission_*λ*_’.

We do wish to point out that our strategy of incorporating a moveable mirror or light deflector into the emission path has also been used for the determination of other parameters, including the faster measurement of fluorescence dynamics [20], determination of the emitter *z*-positions [21], and fluorescence lifetime imaging using standard camera detectors [22].

### Alias-free imaging of single quantum dots

We explored the usefulness of our device via the imaging of single CdSe/ZnS core-shell type quantum dots. This type of emitters has attracted great interest on account of their high brightness, high photostability, and narrow emission spectra [16]. However, they are also well-known for their complex excited-state dynamics, including pervasive fluorescence dynamics over a broad range of timescales [23]. We imaged immobilized quantum dots with emission spectra centered at 655 nm (QDot655) and 585 nm (QDot585) using our device (figure 2a-b), ensuring that the illumination intensities of the lasers were identical such that the ratio of the observed emission spots should reflect the excitation spectra of the quantum dots. The resulting per-lobe intensities aligned with the bulk excitation spectra (figure 2a-b), showing that our Resonator indeed encodes the excitation efficiencies of the fluorophores.

**Figure 2:**
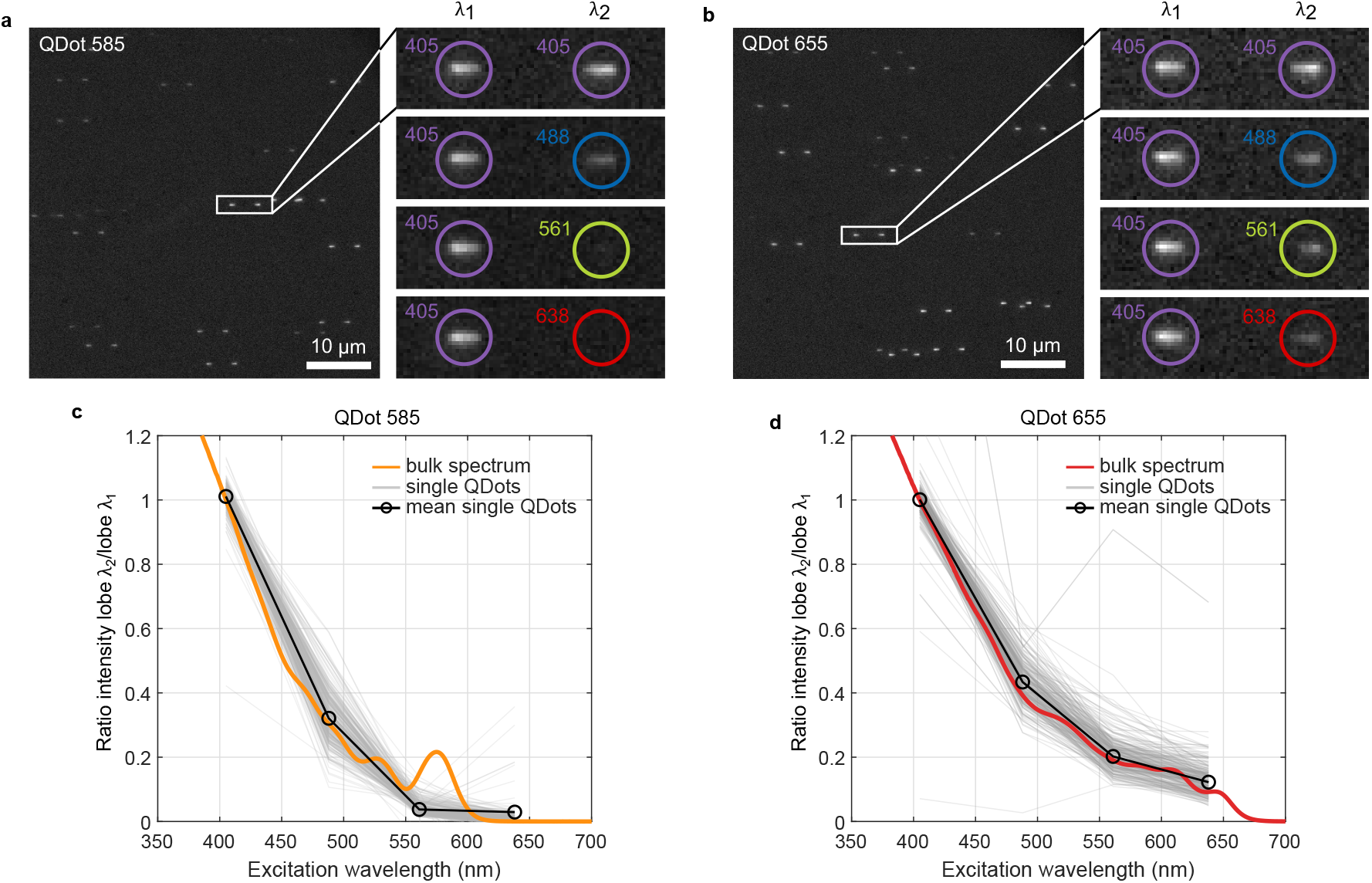
Probing the excitation spectra of single quantum dots. **(a)** Example image obtained on QDot585 emitters, where the inset shows different combinations of the illumination wavelengths. **(b)** Same as panel (a) but for QDot655. **(c)** Fluorescence intensities observed at the different excitation wavelengths normalized to the intensity observed with 405 nm excitation (*n* = 293 quantum dots from 14 fields of view). The colored line shows part of the excitation spectrum provided by the quantum dot supplier. **(d)** Same as panel (c) but for QDot655 (*n* = 231 quantum dots from 14 fields of view).

We next turned our attention to monitoring the fluctuations in the emission of individual quantum dots. Quantum dots are known to display fluorescence intermittencies (‘blinking’) over a large range of timescales [24], which can readily lead to aliasing using sequential multichannel acquisitions. We again immobilized QDot655 emitters on a cover glass and measured their emission over 5000 fluorescence images with a 50 ms exposure time (total duration approximately four minutes), using excitation at 488 and 638 nm. A single image from such a dataset is shown in figure 3a.

**Figure 3:**
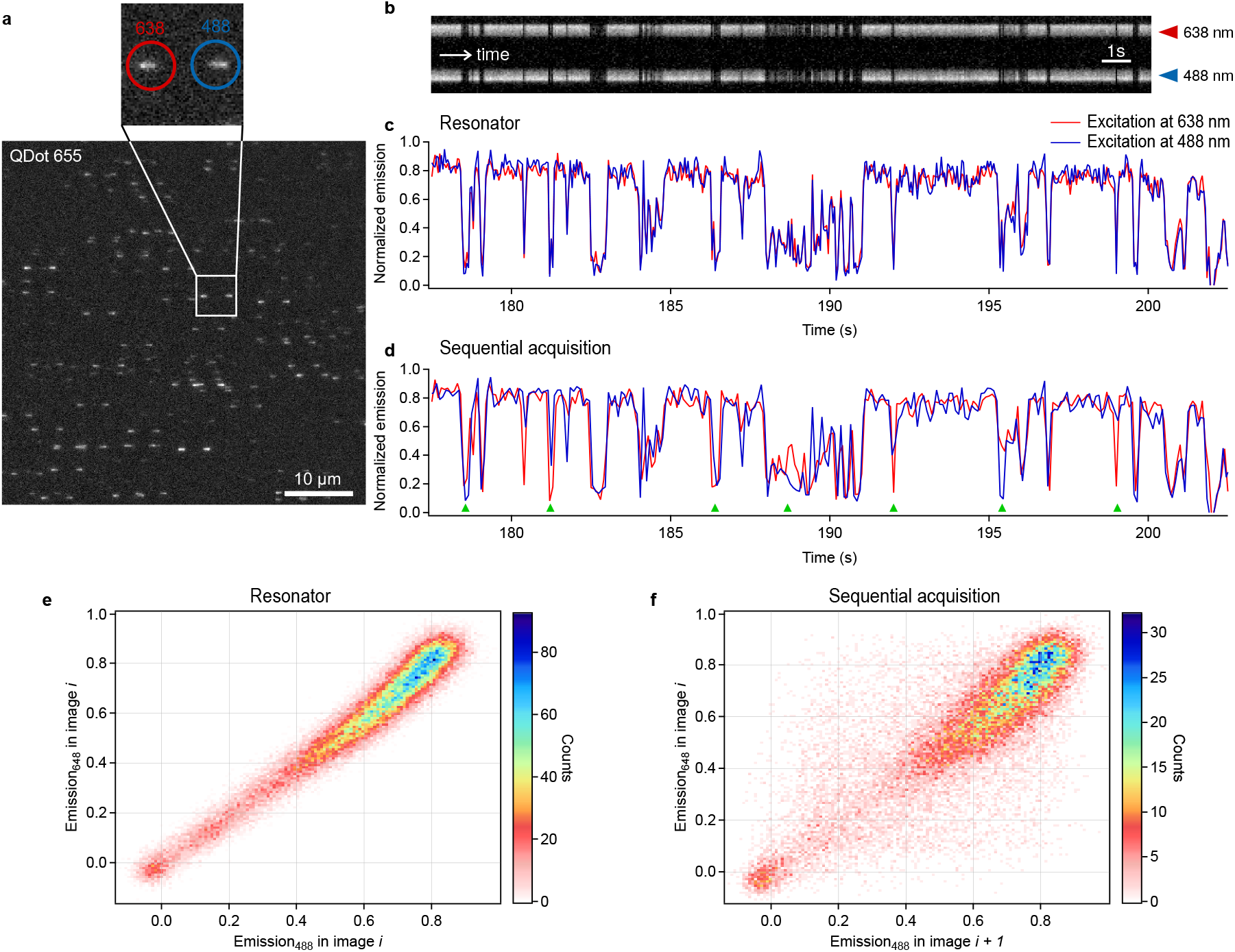
The Resonator suppresses aliasing artifacts in blinking quantum dots. **(a)** Example fluorescence image showing single QDot655 emitters. **(b)** Kymograph excerpt of the quantum dot highlighted with the inset in panel (a). **(c)** Intensity time traces of the quantum dot from panel (a). **(d)** Time traces calculated from those in panel (c) to approximate the effects of classical sequential acquisitions without the Resonator. The green triangles highlight events where artifacts due to temporal aliasing can be observed, though more artifacts can be seen. **(e)** 2D histogram of the fluorescence intensities observed in each image for the Resonator data in panel (c). **(f)** 2D histogram of the fluorescence intensities observed in each image for the calculated sequential acquisition data in panel (d).

Figure 3b shows a kymograph excerpt from an intensity timetrace measured on a single quantum dot, showing the fluorescence intensities observed for each of the excitation wavelengths. Figure 3c shows the same data in a conventional graph. As expected, the traces show pronounced intermittencies on fast timescales. To compare these ‘Resonator’-based acquisitions with conventional sequential acquisitions, we generated a derived dataset that mirrors such sequential acquisitions by taking the intensity from only one of the excitation wavelengths in each image, alternating between the wavelengths in consecutive frames. In other words, we consider only the intensity of the first PSF lobe in image *i* and only the intensity of the second lobe in image *i* + 1, repeating this procedure for the full duration of the acquisition. The resulting intensity timetrace is shown in figure 3d. Comparison of the Resonator-based and sequential timetraces show that sequential acquisitions lead to clear aliasing artifacts, evident in mismatches between the 488 nm and 638 nm excitation intensity traces. Selected instances of this are highlighted using the green markers in figure 3d.

The temporal aliasing seen in sequential acquisitions can suggest the presence of molecular states that do not in fact exist. To illustrate this, we plotted the fluorescence intensity observed upon 638 nm excitation against that observed upon 488 nm excitation, using both the native Resonator data and the sequential-like dataset described above (figure 3e and f). The sequential data shows a main fluorescent population but also shows broadly scattered intensities suggesting the presence of molecular states with shifted excitation spectra that absorb one of the excitation wavelengths more strongly. However, the Resonator data shows that the intensities of the two lobes are highly correlated, indicating that the data can be adequately explained as blinking involving a single fluorescent state with a constant excitation spectrum. These observations readily demonstrate the value of our device for the distortion-free measurement of fast processes.

### Simultaneous excitation and emission spectroscopy of quantum dots

We next sought to incorporate both excitation and emission information into the measurement. To do so, we placed a 630 nm long pass dichroic mirror in close proximity to the resonant scanning mirror, as shown in figure 4a. This configuration creates a PSF with 3 spots, where the relative intensities of the spots depend on both the emission and excitation spectra of the emitter: fluorescence photons with emission above 630 nm are transmitted by the dichroic, creating two emission spots as previously described, while photons with a wavelength below 630 nm are reflected by the dichroic and create a third emission spot regardless of the excitation wavelength that gave rise to this emission. Both emission bands are further separated by the presence of optical filters that block the 638 nm laser illumination.

**Figure 4:**
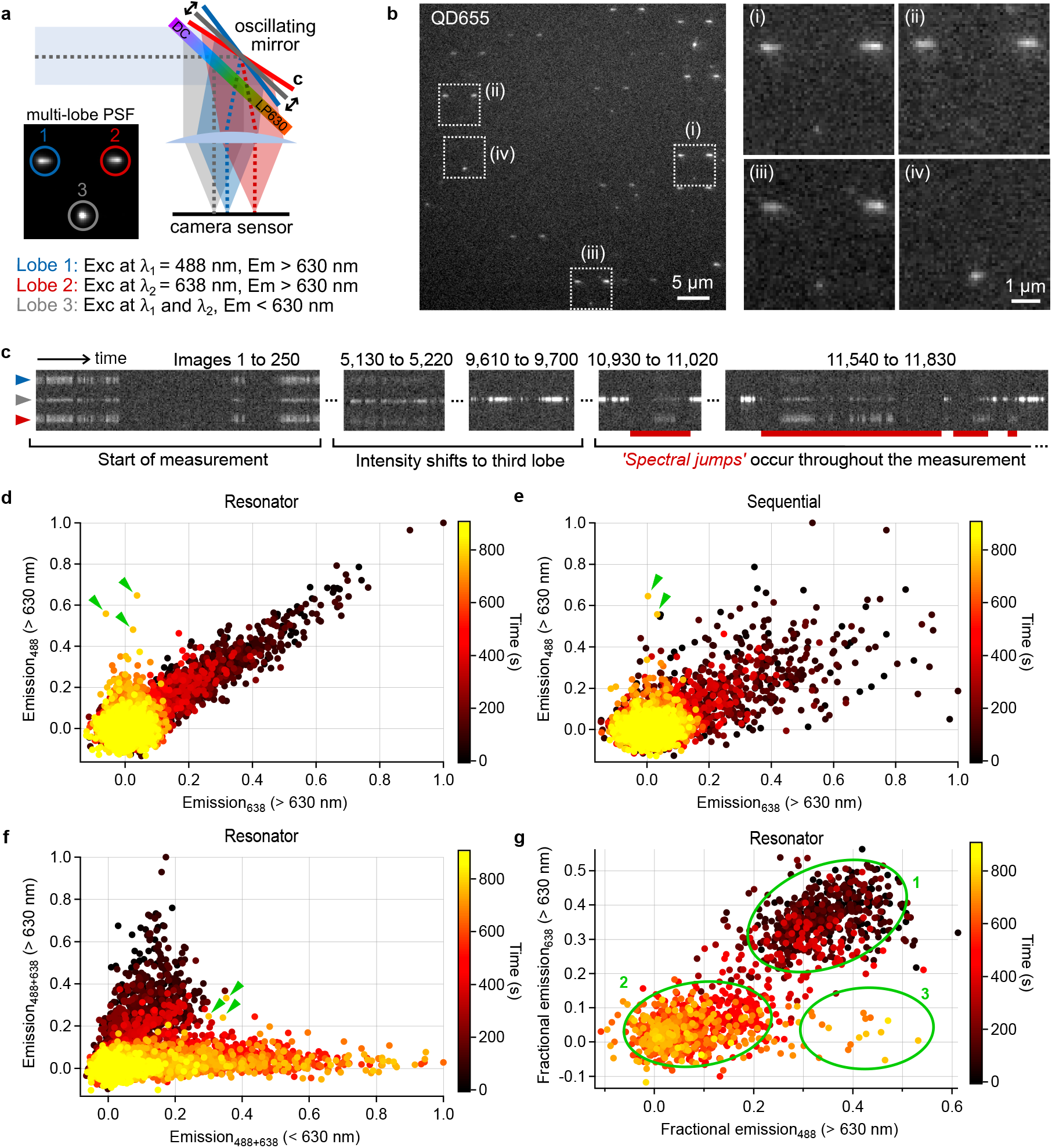
Combined single quantum dot excitation and emission spectroscopy. **(a)** Schematic illustration of the three-lobe optical setup. **(b)** Example image of quantum dots (QD655) on a cover glass, with 4 insets showing varying intensity distributions indicating spectral differences. **(c)** Kymogram showing spectral dynamics in a single QDot655 emitter. A limited set of intervals of interest has been selected. **(d)** Temporally colour-coded scatter plot of the intensities observed in the excitation lobes emitting above 630 nm. The green markers in this and following panels are discussed in the main text. **(e)** Calculated sequential acquisition data approximating what would be observed for the data in panel (d) without the Resonator. **(f)** Temporally color-coded scatter plot distribution of the emission intensities above and below 630 nm. **(g)** Temporally color-coded scatter plot of showing the fractional contribution of the indicated emission to the total emission (sum of the three lobe intensities) observed in each image. To enhance the visual clarity, only images in which the total emissionwas over a threshold were included in this visualization, as is discussed in more detail in the materials and methods section.

Figure 4b shows an example fluorescence image and expansions acquired on QDot665 emitters, showing a high spectral diversity in the emission of the individual quantum dots. This diversity was evident also in the individual quantum dot timetraces, an example kymograph of which is shown in figure 4c. In addition to undergoing transitions in excitation and emission spectra, the quantum dots where also observed to undergo a blueshift upon irradiation, which has been widely described as ‘spectral blueing’. This effect has been attributed to a decrease in quantum dot size due to photo-oxidation [25–27] with a rate that depends on the excitation wavelength [28].

Figure 4d analyzes the spectral diversity in more detail by showing the relative intensities of the 488 and 648 nm-excited fluorescence emitted over 630 nm for this particular quantum dot, taking advantage of our Resonator. Most of the data points appear to lie on a single line, indicating the presence of dynamics involving a fluorescent state with constant excitation efficiences for both excitation wavelengths. Figure 4e, in contrast, approximates what would be observed using a classical sequential acquisition, revealing a more scattered distribution of the intensities that is much less straightforward to interpret. The intensity of the third PSF lobe provides information on the emission spectrum of the quantum dot, confirming a gradual blueing of the emission towards shorter wavelengths (figure 4f).

Closer inspection of figure 4d shows the presence of several data points that do not align with the overall trend, indicated with green markers on the graph. These points highlight the presence of a third state characterized by a higher efficiency for 488 nm excitation compared to 648 nm excitation. While these points can also be observed in the sequential data, they are obscured by the scattering induced by the temporal aliasing (figure 4e). Figure 4f additionally reveals that this third state has an emission spectrum in between the two detected wavelength bands. However, this data does not easily reveal whether additional points may also have arisen from this state but are intermingled with observations made on other states. To enhance the discriminatory power of this analysis, we visualized information from all three PSF lobes at once by graphing the fraction of the emission that the Resonator lobes contribute to the total detected quantum dot fluorescence in each image (figure 4g). In this figure, three different states can be clearly discerned, with state 1 the initial state, state 2 the state obtained upon emission blueing, and state 3 the intermediate state that is transiently formed in the measurement. This finding highlights the additional information enabled by our device.

### Ratiometric Ca^2+^ sensing in firing neurons

The previous experiments used small scanning angles of the resonant mirror in order to encode the excitation information into the fluorophore PSF. Our device can also be used in a more classical image splitting arrangement by increasing the amplitude of the resonant scanner movement combined with the introduction of a field stop. The responses to the different excitation wavelengths then appear as distinct regions on the camera sensor, essentially mirroring the use of a conventional image splitter for emission multiplexing (figure 5a). One area where such a system finds obvious use is in the imaging of biosensors, fluorophores for which the emission is conditional on a well-defined aspect of their environment [29]. Within the large biosensor family, ratiometric biosensors encode this information as a change in the ratio of two excitation and/or emission bands. We chose to focus on the ratiometric Ca^2+^ biosensor FuraRed [12],which encodes the local calcium concentration into the ratio of the fluorescence intensities excited at 405 nm and 488 nm. The fast dynamics of neuronal signaling have given rise to multiple strategies to deliver faster excitation-ratiometric measurements (e.g. [30]).

**Figure 5:**
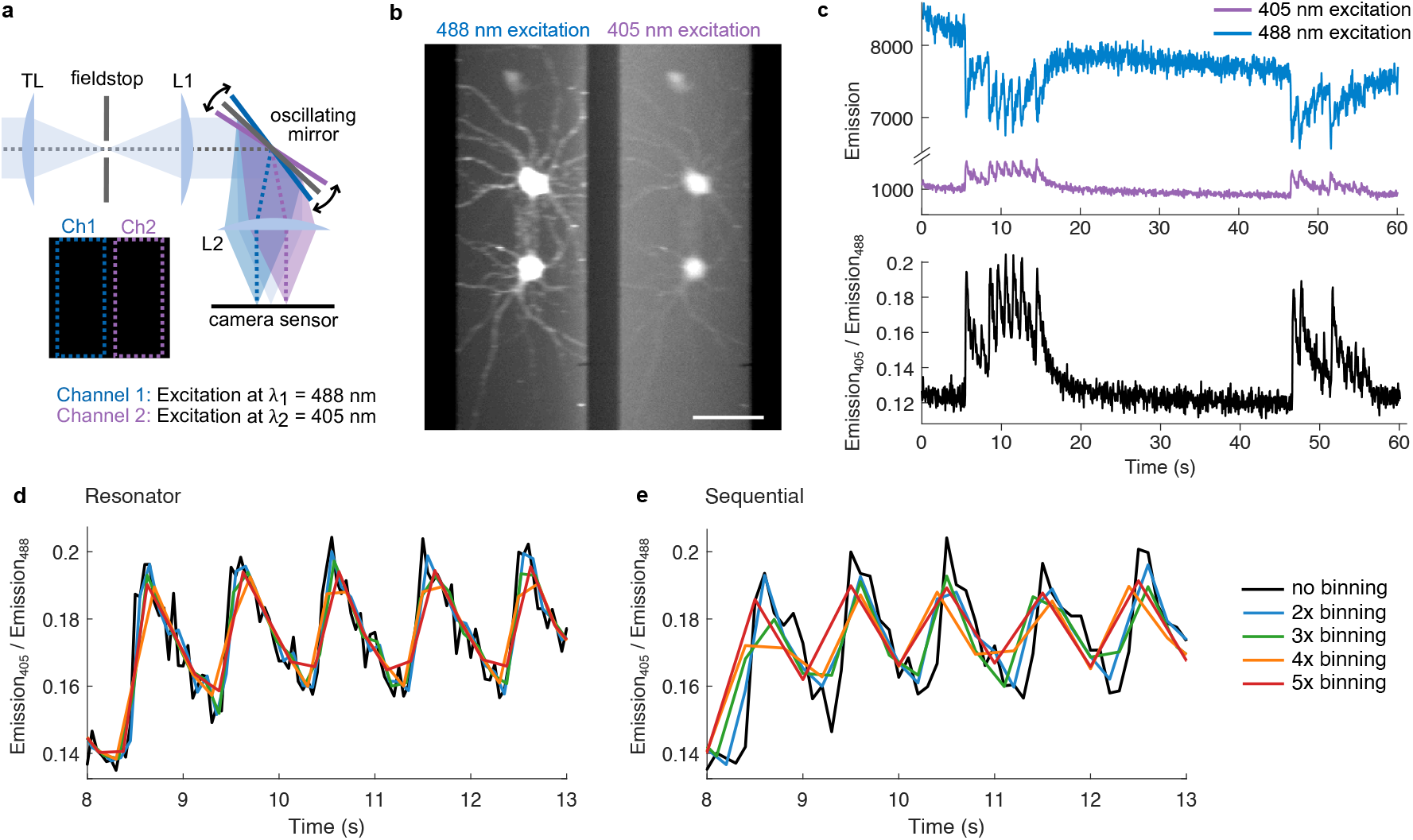
Ratiometric calcium imaging in firing hippocampal neurons. **(a)** Schematic illustration of the optical layout. **(b)** Average image calculated from all 1560 fluorescence images acquired on two neurons, showing the two excitation channels. The color lookup table is different for the 405 and 488 nm image so that both channels are similarly observable.**(c)** Integrated intensities observed for the upper cell from panel (b) as the ratio of the intensities. **(d)** Calculation of the excitation ratios for different temporal binning values. **(e)** Same as panel (e) but for calculated sequential acquisitions.

We applied the Resonator to the imaging of Ca^2+^ dynamics in firing mouse hippocampal neurons stained with FuraRed, where the different excitation channels are imaged onto distinct regions of the image sensor (figure 5b). The action potentials are clearly visible from the intensity traces observed at each excitation wavelength as well as the ratio between them (figure 5c). We again compared our Resonator-based acquisition with a derived dataset showing what would be observed using sequential acquisitions, additionally varying the temporal resolution by binning adjacent images together. The calcium responses in these measurements were slow enough that comparatively little temporal aliasing arose even in sequential measurements, reflecting also that our microscope did not possess an incubator and the measurements were accordingly performed at room temperature. However, comparison of the Resonator-based and sequential measurements for binned data (figure 5d and e) clearly demonstrates that our device can indeed eliminate aliasing artifacts also under these conditions, showing that this strategy can also be applied to samples that are too dense to support PSF-encoding of the information or where such encoding is undesirable.

## Conclusion

In this work we have presented a strategy for near-simultaneous excitation multiplexing based on synchronizing the excitation light with fast displacements of the sample image on a camera sensor. We developed a specific implementation called the Resonator, which can encode two different excitation wavelengths within a single fluorescence acquisition by introducing a rapidly oscillating mirror into the emission beam path, such that each fluorophore can be observed as two distinct emission spots corresponding to each of the excitation wavelengths. The distance between these spots can be tuned by changing the amplitude of the mirror motion, where small amplitudes make it possible to use the device as a type of PSF engineering, while larger amplitudes and the introduction of a field stop allows more conventional image splitter-like operation. Further excitation wavelengths could be introduced by synchronizing their illumination with different ranges of the scanner motion. The resulting information can be used to distinguish different fluorophores and/or to gain more information on their molecular state and local environment.

The crucial advantage of our device arises because common resonant scanners readily provide scanning frequencies up to 16 kHz or even faster, allowing the different excitation wavelengths to be probed within just a few tens of microseconds using comparatively inexpensive equipment. Compared to the per-image sequential acquisition usually used for excitation multiplexing, our method allows much faster sampling and eliminates (much of) the aliasing artifacts that otherwise lead to distortions in the measurement, as we demonstrated in this study using the imaging of quantum dots and hippocampal neurons. While this implementation has made use of a resonant scanner, our approach could also be combined with alternative fast scanning mirrors such as MEMS-based devices. Beyond the applications shown here, the device could also be applied to e.g. correct for acceptor blinking in single-molecule FRET, or to identify different emitters in methodologies such as single-molecule localization microscopy.

Our approach can also be combined with other functionalities to achieve additional information content. One example is the introduction of an additional, stationary, dichroic mirror that results in a three-lobe PSF, as we have demonstrated here. The Resonator and similar systems could also be combined with conventional dichroic-mirror based image splitters, simultaneously delivering information on both the excitation and emission spectra. The PSF engineering of the Resonator could also be combined with other PSF-engineering modalities such as the introduction of astigmatism or more complex phase manipulations to encode *z*-positions or emission spectra of the fluorophores, since our strategy uses only a flat reflective surface and therefore does not perturb any other phase manipulations that may have been introduced.

Very recently, a related approach was described in which a fast scanning element was used to create three different subimages on a camera sensor [31]. Our work distinguishes itself from this work by offering a simpler optical layout that additionally enables the PSF-engineering approach described here, as well as the use of an additional dichroic to also encode emission information.

In conclusion, we have developed a strategy and optical device that deliver near-simultaneous excitation multiplexing in any situation where fast processes must be monitored by observing the fluorescence emitted in response to different excitation wavelengths. By providing (nearly) alias-free multiplexing based on excitation wavelength, our device readily expands the available strategies for multispectral fluorescence imaging and spectroscopy.

## Materials and methods

### Resonator optical setup

An imaging system was built based on an inverted microscope body (Eclipse Ti2-E, Nikon). An L6Cc laser combiner (Oxxius) with 5 laser lines (405 nm, 445 nm, 448 nm, 561 nm, 638 nm) was used for excitation. The laser beams were focused to a spot in the back focal plane of a Plan Apo 100 × 1.49 NA oil-immersion objective (Nikon). The excitation light and fluorescence were filtered and split using a quadband filter set with an excitation filter (ZET405/488/561/640xv2, Chroma), dichroic beamsplitter (ZT405/488/561/640rpcv2, Chroma) and emission filter (ZET405/488/561/640m, Chroma). An EMCCD camera (ImageEM EMCCD C1900-13, Hamamatsu) was used for imaging.

For the 2-lobe optical setup (figure 1a), two lenses (L1, AC254-080-A; L2, AC254-050-A, Thorlabs) were placed in a 4f configuration and the mirror of the resonant scanner (RS, S/N46970 SC30-1KHz, 20 × 15 cm, Electro-Optical Products Corp.; with driver PLD-2S, Electro-Optical Products Corp.) was placed in the conjugate back focal plane of the objective. The resulting virtual pixel size of the system is 104 nm × 104 nm.

For the 3-lobe optical setup, a dichroic mirror (DMLP630T, Thorlabs) is placed in front of the mirror of the resonant scanner as depicted in figure 4a. The reflective surface of the dichroic and mirror of the resonant scanner were placed as close together as possible to limit the optical path difference between light that is reflected by the dichroic and reflected by the mirror of the resonant scanner. The dichroic is mounted in a kinematic mount (KM100C, Thorlabs) to allow fine positioning of the PSF-lobe that is reflected by the dichroic with respect to the lobes reflected by the resonant scanner.

Custom electronics were used to synchronize the laser excitation with the mirror position. Custom software was used for image acquisition.

### Quantum dot sample preparation and measurements

Cover glasses were plasma cleaned for 45 minutes and rinsed with MilliQ water. The quantum dot working solution was prepared by centrifuging the stock solution (QD655, cat. no. Q11621MP, and QD585, cat. no. Q11411MP, Invitrogen) for 3 min at high speed to remove aggreggates. 1 µL of the supernatant was then diluted in 1 mL MilliQ, and 50 µL of a further 1:200 dilution was deposited directly on the cover glass. The quantum dots were left to settle on the surface for 30 minutes before drying.

A camera exposure time of 50 ms and a duty cycle of the illumination of approximately 26% was used (*i*.*e*. during each mirror oscillation, the sample if illuminated by each laser for 128-132 µs as measured from the width of the electrical signal used to trigger the light sources).

For the experiment shown in figure 2, the power of the 405 nm, 488 nm, 561 nm and 638 nm lasers were matched by measuring the laser powers after the objective, such that the relative intensities of the PSF lobes should indicate the absorption of the emitter at the respective wavelengths.

### Quantum dot data analysis

Quantum dots were identified and localised as follows: an average projection of the image stack is generated on which localisation will be performed. The average projection is background corrected by subtracting a spatially median filtered (50×50 pixel kernel) temporal median intensity projection of the dataset. To identify possible quantum dots, a maxima search is performed on the background corrected average intensity projection filtered using a difference of Gaussian filter. An asymmetric Gaussian is then fitted to all the candidates.

The different emission spots were then assigned to individual emitters based on the knowledge of the expected PSF, which is fully determined by the instrumental settings. In the case of three-lobe PSFs, the emission spot created by reflection from the dichroic could be further discerned because it is slightly more symmetric (ratio of *σ*_*x*_ and *σ*_*y*_ from the asymmetric Gaussian fit closer to 1) compared to the spots created by the moving scanning mirror. Emission spots that were too close to the image border or could not be connected unambiguously within a single PSF were rejected. The resulting list of emitter coordinates was then used to create intensity timetraces for each individual lobe by summing the pixel values in a small neighborhood (7 by 7 pixels). In figure 4g, only datapoints for which the total fluorescence (sum of all three lobes) exceeds 22.6% of the maximal observed total fluorescence were used.

### Cell culture mouse hippocampal neurons

1-2 day old C57BL/6J mouse pups were quickly decapitated before dissection. Hippocampi were dissected in Sylgard dishes containing cold sterile Hank’s buffered salt solution (HBSS in mM: 5.33 KCl, 0.44 KH2PO4, 137.93 NaCl, 0.34Na2HPO40.7H2O, 5.56 D-glucose and 10 Hepes). The tissue was incubated in 0.25% trypsin-ethylenediaminetetraacetic acid (EDTA) (Gibco) supplemented with 200 U/ml DNase (Roche) for 10 min at 37µ°C. After three consecutive wash steps with washing buffer (neurobasal medium (Invitrogen) supplemented with 1.59 mg/ml BSA, 0.5% penicillin/streptomycin, 5 mg/ml glucose, 5.5 mg/ml sodium pyruvate, and 200 U/ml DNAaseI) the tissue was mechanically dissociated by trituration. After centrifugation, cells were resuspended and plated on 18 mm diameter coverslips, coated with poly-D-Lysine, suspended over a glial support layer using paraffin dots. The glial support layer was derived using an identical protocol on midbrain and cortical tissue harvest from the same pup. Primary hippocampal neuron-glia co-cultures were grown in a 37µ°C, 5% CO2 incubator in Neurobasal-A media (Thermo Fisher Scientific) supplemented with 0.5% penicillin/streptomycin (Lonza), 2% B27 (Gibco), 0.02 mg/ml insulin (Sigma), 50 ng/ml nerve growth factor (Alomone Labs), and 0.5 mM Glutamax (Thermo Fisher Scientific). Half of the media volume was replaced every 3 days. Imaging was performed after 7-10 days in vitro. All procedures were approved by the Animal Ethics Committee of the KU Leuven.

### Calcium imaging and stimulation of neurons

Coverslips were loaded with 1 µM FuraRed-AM in HEPES buffered solution containing in mM (140 NaCl, 5 KCl, 10 HEPES, 2 CaCl2, 2 MgCl2 and 10 D-glucose, adjusted to pH = 7.4 using NaOH) for 20 min. After 5 min wash with HEPES buffer, the coverslips were transferred to a custom built holder and mounted on the microscope stage. A custom-built miniature bipolar electrode was carefully positioned over the field of view using a micromanipulator. FuraRed was excited by 405 nm and 488 nm laserlight, with respectively 40 ms and 30 ms exposure times. At a set times during the recording, electrical field stimulation was applied via the bipolar electrode that was connected to a high current stimulus isolator (WPI, A385) and Master 8 (A.P.I) to generate the desired current amplitude (15 mA) and stimulus paradigm (10x, 1ms, 1Hz) respectively.

### Analysis of calcium signalling data

Intensity traces were generated by integrating the fluorescence over the entire cell body, using a mask defined as all points exceeding a treshold in an image consisting of the average of all acquired fluorescence images (6018 and 3868 counts for 488 nm and 405 nm channels respectively for the data in figure 5b). A constant background signal was removed based on the average value of a 10×10 pixel region devoid of cells (1718 counts for the data in figure 5b). The ‘ratio 405/488’ trace from was obtained by calculating the ratio of the extracted 405 nm and 488 nm intensity traces at each time point. The binned intensity trajectories were obtained by adding consecutive data points of the original 405 and 488 nm intensity traces.

## Acknowledgements

We thank Steven F. Lee (University of Cambridge) for discussions and support. R.V.D.E. thanks the Research Foundation-Flanders (FWO Vlaanderen) for a doctoral fellowship. M.M. was supported by a Marie Skłodowska-Curie postdoctoral fellowship. This work was supported by the European Research Council through grant 714688 NanoCellActivity and the Research Foundation-Flanders through grant G090819N.

## Author contributions

W.V., P.D., and M.M. designed research. P.D. supervised research. R.V.D.E., E.B., M.M., B.A., and W.V. constructed the optical device. E.B., R.V.D.E., T.V., and P.V.B. prepared samples. E.B., R.V.D.E., M.M, T.V., and P.V.B. performed experiments. E.B., W.V., and P.D. analyzed data. P.D. and E.B. wrote the manuscript with input from R.V.D.E. and W.V.

